# Ximmer: A System for Improving Accuracy and Consistency of CNV Calling from Exome Data

**DOI:** 10.1101/260927

**Authors:** Simon P Sadedin, Justine A Ellis, Seth L Masters, Alicia Oshlack

## Abstract

Detection of copy number variation (CNVs) is a challenging but highly valuable application of exome and targeted high throughput sequencing (HTS) data. While there are dozens of CNV detection methods available, using these methods remains challenging due to variable accuracy both across different data sets and within the same data set with different methods. We propose that extracting good results from CNV detection on HTS data requires a systematic approach involving rigorous quality control, adjustment of method parameters and calibration of confidence measures for filtering results. We present Ximmer, a tool which supports an end to end process for applying these procedures including a simulation framework, CNV detection analysis pipeline, and a visualisation and curation tool which enables interactive exploration of CNV results. We apply Ximmer to perform a comprehensive evaluation of CNV detection on four data sets using four different detection methods, representing one of the most comprehensive evaluations to date. Ximmer is open source and freely available at http://ximmer.org (example results are viewable at http://example.ximmer.org).

## Background

In recent years, high throughput sequencing (HTS) of DNA has become an essential tool in biomedical science with a vast range of applications spanning both clinical and research investigations. In clinical settings, whole exome sequencing (WES) and custom targeted gene panels are especially important and have enabled significant improvements in the rate of diagnosis for genetically heterogeneous disorders [1]. WES has also had a profound impact on disease research, by allowing researchers to comprehensively search for protein altering genetic variation. As a result of these advances, the rate of discovery of new Mendelian disease genes has seen substantial improvements in recent years (X. Zhang 2014).

While WES has proven highly effective, this success has been based predominantly on the detection of single nucleotide variants (SNVs) and small insertions and deletions (indels). Larger variants, such as copy number variants (CNVs), are not routinely ascertained from WES data. Nonetheless, CNVs are frequently disease causing, both as the primary genetic lesion for disorders such as α-thalassemia (Stankiewicz and Lupski 2010), Charcot-Marie-Tooth neuropathy and Smith-Magenis-Syndrome, as well as a rare cause for a wide range of mendelian diseases. In particular, single copy deletions can be pathogenic for any disorder caused by haploinsufficiency. To detect CNVs, patients are often screened for CNVs using SNP or array-CGH microarrays prior to use of WES. However, affordable microarrays have limited resolution and add time, cost and complexity to the overall diagnostic workflow. There are consequently significant potential advantages if CNVs can be ascertained directly from WES.

CNVs can be detected from three primary signals in HTS data. These are: anomalous mapping of paired end reads that span CNV breakpoints (PE signals), the splitting of individual reads by CNV breakpoints (SR signals), and fluctuation in the coverage of reads falling in the body of a CNV (the read depth, or RD signal). While all of these signals are effective in whole genome data, the breakpoints of CNVs usually fall between the regions targeted by WES. Therefore only the RD signal is reliably observable in WES data. The RD signal has been shown to be informative due to a strong correlation of copy number with read coverage depth [2]. However, detection of CNVs is confounded by a range of other factors that also influence read coverage depth. Therefore, these factors must be corrected, and failure to do so can result in significantly degraded accuracy.

Numerous methods have been developed to detect CNVs based on the RD signal. Examples include ExomeDepth [3], ExomeCopy [4], XHMM [5], cn.MOPS [6], ExomeCNV [2], CoNVEX[7], EXCAVATOR [8], CoNIFER [9], CANOES [10], CODEX [11], and many others. The authors of these tools have often cited high sensitivity and specificity for their methods. However, independent comparisons frequently fail to replicate their findings. For example, Guo et al. reported ExomeDepth having sensitivity of only 19% [12], while Ligt et al. observed a sensitivity of 35% [13]. In the same studies, sensitivity of CoNIFER was cited as 3% and 29% respectively, compared to the original evaluation estimate of 76%. In some contexts, high accuracy is reported. For example, Jo et al [14], Ellingford et al [15] and Feng et al [16] all cited 100% sensitivity and high specificity for detection of larger CNVs encountered clinically, in each case using high coverage data. However, the circumstances in which high accuracy can be achieved are currently not well understood.

Recent studies have compared performance across multiple data sets [17–19], highlighting the problem of variability in the performance of CNV calling as well as high false positive rates [20]. Some of the performance variability observed in these studies may be due to differences between the data sets and sequencing design such as the number of samples, read length, insert size, and mean read depth. Also of critical importance is the size and type of CNVs assessed. However, even when these known technical factors are controlled, significant variability is often still observed between data sets.

In this work we present Ximmer, a software tool that improves CNV calling reliability by enabling users of CNV detection tools to efficiently assess and tune performance. Ximmer contains three parts: a simulation method, an analysis pipeline, and a graphical report. First, Ximmer simulates synthetic single copy deletions in existing WES data. Then, the analysis pipeline automates detection of the deletions with up to 5 commonly used CNV detection methods. Finally, the graphical report shows the combined CNV calling results, including a suite of plots that give insight into the accuracy achieved and strategies for improving performance.

In this article we explain the implementation details of Ximmer, and demonstrate how using Ximmer improves CNV detection accuracy. We show results from four CNV callers on four datasets representing different exome capture kits and different sequencing depths. Our results concur with previous studies, finding that CNV detection performance is highly variable with within and between data sets. However, we show that using Ximmer to gain insight into the variability enables optimisation of the CNV calling, and improves detection of real CNVs. Ximmer offers an integrated framework that is easy to use and freely accessible, from http://ximmer.org. An example of Ximmer output is available at http://example.ximmer.org.

## Methods

The Ximmer process consists of a series of steps designed to optimise CNV detection performance. The steps consist of: (i) simulation of CNVs in the user’s data, (ii) execution of CNV callers to find both real and simulated CNVs, (iii) quality and accuracy assessment to discover optimal settings for CNV calling, and finally, (iv) filtering of results to produce a curated CNV list. This process can be time consuming if conducted manually, however Ximmer automates all of the steps needed. The high level process is depicted in Figure 1.

**Figure 1:**
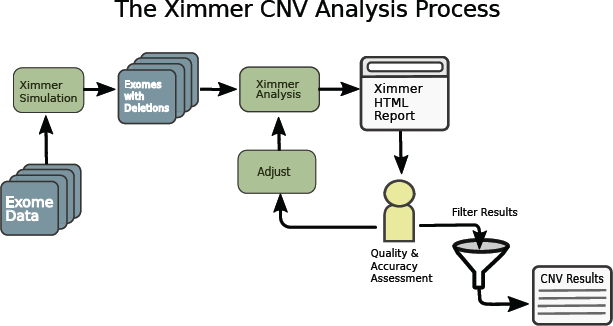
The Ximmer Process - Ximmer consists of three high level steps. In the first step, simulated CNVs are added to a set of sequence alignments in BAM or CRAM format. This creates new BAM files containing simulated CNVs which are passed to the integrated analysis pipeline. The analysis pipeline runs up to 5 different CNV detection methods and collates the results into a graphical report which generates insight into the performance of the tools and possible avenues for improvement. Finally, when the analysis is optimised provides an interface to filter CNVs, review and interpret them using the built in CNV curation tool.

## Simulation

Simulation is the first and most important element of the Ximmer method. By simulating CNVs in the user’s own data, Ximmer generates both a prediction of the CNV calling performance, and also insights regarding how to improve performance. To simulate CNVs, Ximmer takes advantage of the exclusive use of the RD signal by WES based CNV detection methods. Specifically, Ximmer removes reads that overlap selected target regions, such that the RD signal is reduced to match the predicted level associated with a single copy deletion. Ximmer focuses on deletions because depleting reads is significantly simpler than realistically synthesising and adding new reads. While the approach is strictly limited to deletions, the inferences derived are still likely to apply to other CNV states, because in most CNV calling tools the same underlying statistical principles are applied regardless of the number of copies being searched for.

The simulation process begins by randomly selecting the genomic region to become the deletion “target”. The reads mapping to these locations can then be depleted using two alternate methods, referred to as “Downsampling” and “X-Replacement”. The downsampling method randomly removes each read overlapping the deletion target with probability of 0.5, based on an assumption that the relationship between copy number and read depth is linear. By contrast, the X-Replacement method avoids this assumption. The X-Replacement method replaces reads mapping to X chromosome deletion target regions in a female sample with an adjusted number of reads from the same genomic regions in a male sample. This method exploits the true difference in copy number between male and female X chromosomes to avoid the assumption of linearity implied by downsampling. The X-Replacement method also ensures that other aspects of the reads are preserved in a realistic manner, such as the zygosity and phasing of overlapping SNVs and indels. Further details of the simulation implementation are provided in the supplementary methods (S-1). The result of the simulation step is a new set of alignments (BAM files) for the whole exome, but with deletions simulated in selected regions.

## CNV Analysis Pipeline

The second step in the Ximmer process is to analyse the data containing simulated CNVs to produce CNV calls. Ximmer provides a built in analysis pipeline that automatically installs, configures and runs 5 commonly used CNV detection methods. These tools are: ExomeDepth, XHMM, cn.MOPS, CoNIFER and CODEX. The analysis pipeline is constructed using Bpipe [21], a framework for creating bioinformatic workflows. In addition to running the CNV detection tools, Ximmer performs any necessary pre-processing required by the tools and also post processes the results to merge and annotate the resulting CNV calls. Additional CNV callers can be added to Ximmer with only a small effort through the extensible Bpipe framework.

The analysis produces a report in HTML format that contains a full summary of all the simulated deletions, along with a range of plots and tables to highlight CNV calling performance and potential quality issues.

## Results Assessment

Once CNV analysis has been performed, the next step in the method is to critically review the HTML report to assess performance of the CNV callers for detecting the simulated deletions, and to evaluate options for improving the results.

### Quality Assessment

Three plots are of particular relevance in understanding potential quality issues. These are the Sample Counts, Genome Distribution and Quality Score Calibration plots.

The Sample Counts plot (Figure 2A) shows the distribution of the number of CNV calls among the samples, separately for each CNV caller. In most studies we expect the number of CNV calls to be similar for each sample. If some samples contain a disproportionate fraction of the total CNV calls, it is likely that there is a problem with the sample quality. It may be desirable to either remove the samples from the CNV calling altogether, to adjust the caller settings to compensate, or to isolate poor quality samples from use in normalising other samples.

**Figure 2:**
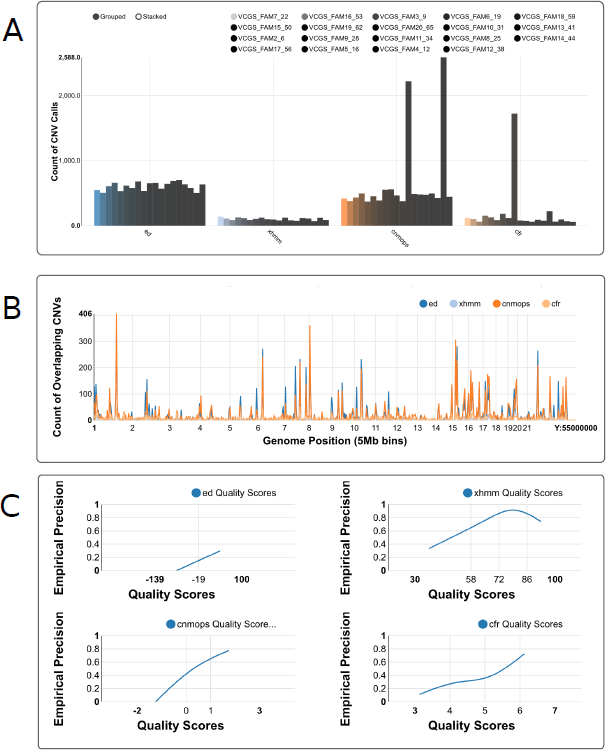
Screen shots of Ximmer Quality and Accuracy Plots. **A.** Sample counts plot, showing the number of CNV calls for each sample, by each CNV caller. **B.** Genome distribution plot, showing frequency of CNV calls along the genome. **C.** Quality score calibration plot, showing relationship of empirical precision to quality score.

The Genome Distribution plot (Figure 2B) divides the genome into 5 megabase bins and displays the number of CNVs overlapping each bin. Clicking on a particular region displays an enlarged plot encompassing that region for more detailed inspection. If particular regions contain very large numbers of CNV calls, it may be desirable to remove these from the target regions used for calling, as their presence may distort quality statistics and degrade overall calling accuracy.

The Quality Score Calibration plot (Figure 2C) assists in interpreting the confidence measures (or quality scores) assigned to CNVs by the CNV callers. For each caller, Ximmer groups the whole CNV call set into approximately 5 quality score bins that collectively span the full range of values assigned by the caller. Ximmer then calculates the fraction of calls categorised as true positives in each bin as an estimate of the precision. The estimates are plotted as a line to illustrate the empirical relationship between precision and quality score for each CNV caller. When quality scores are well behaved it is expected that the precision should increase monotonically as quality score increases. Failure to observe this relationship suggests the caller may produce high confidence false positives, in which case filtering by quality score alone may be insufficient to reduce the false positive rate. As with the Sample Counts plot, it may be appropriate to review normalisation settings for methods if quality scores assigned by tools are not well behaved.

### Accuracy Assessment

After reviewing the quality assessment the next step in the Ximmer process is to review the accuracy estimate. This is presented using a plot designed to mimic a traditional “Receiver Operator Characteristic” curve, but displayed using absolute measures to better accommodate the unknown positions of true negatives in CNV calling. Instead, the ROC-style plots show how the detection of simulated true positives (Y-axis) changes with the number of detections not part of the simulation (false positives, X-axis) as results are progressively filtered to lower significance levels. It should be noted that false positives are defined as regions that are not simulated to be deletions but they could actually be true positives from the sample itself. Unlike comparisons of absolute sensitivity and precision, this method primarily compares the ranking of true and false positives, and thus takes into account the utility of confidence measures output by tools for filtering the results. For the CNV calling tools included in Ximmer, the confidence measure used for ranking results was chosen in each case by consulting the documentation or by discussion with the tool author (Supplementary Methods, Table S1).

The initial display of the ROC-style curve shows the accuracy for the whole set of simulated deletions. As a first step this can suggest an appropriate level at which to filter results so that the optimal level of sensitivity and specificity is achieved. However, the plot can be interactively adjusted, to show performance of a subset of CNVs within specific size ranges.

### CNV Discovery

Once the performance of the CNV callers is well understood, the final step in the Ximmer process is to filter the CNV calls according to the decided quality filtering thresholds. The remaining CNVs may then be manually reviewed in Ximmer’s CNV curation interface. The interface includes a range of annotations to support interpretation of the likelihood that a CNV is real, and whether the CNV is potentially of functional interest. The annotations include overlapping genes, population frequency of relevant CNVs from the Database of Genomic Variants (DGV [22]), overlapping single nucleotide variants (SNVs) or indels, and a pictorial diagram of the read depth deviation over the CNV region.

If desired, the discovery of real CNVs can be from the same analysis result set containing simulated CNVs. This approach relies on an assumption that simulated and real CNVs of interest are unlikely to overlap. Alternatively, Ximmer can be re-executed on the original raw data with simulation disabled to derive a stand-alone result set.

## Data Sets

To demonstrate the application of Ximmer, we applied it to four data sets representing different Illumina sequencing platforms, exome captures, read configurations and sequencing depths (Table 1).

**Table 1:** Data sets analysed with Ximmer.

| Capture | Samples | Capture Size (Mb) | Read Length | Mean Read Depth |
| --- | --- | --- | --- | --- |
| SureSelect v5 | 16 Male,<br>14 Female | 51.2 Mb | 2 x 100 | 30 |
| Nextera 1.2 | 24 Female,<br>28 Male | 45.3Mb | 2 x 100 | 120 |
| Nimblegen v2 | 19 Female,<br>19 Male | 47Mb | 2 x 75 | 60 |
| TruSeq / Custom<br>Broad Capture | 19 Female<br>16 Male | 37.5Mb | 2 x 150 | 90 |

The SureSelect data set was produced as part of an unrelated research program, the Nextera data was created as part of the Melbourne Genomics Health Alliance demonstration project (https://www.melbournegenomics.org.au/) and the TruSeq data was created by the Broad Institute, Center for Mendelian Genomics. The NimbleGen data set was downloaded from the Sequence Read Archive (SRA) from a previous study as part of the Simons Foundation Research Autism Initiative (SRP010920, [23]).

The SureSelect, Nextera and Nimblegen data sets were analysed in house to produce alignment files in BAM format using Cpipe (Sadedin et al. 2015). The TruSeq data set was produced and analysed at the Broad Institute using the institute’s standard analysis pipeline, also based on GATK.

## Results

### Ximmer Simulations

In order to demonstrate Ximmer we applied it to four different exome datasets with a variety of different properties (Table 2). We configured Ximmer to simulate between 2 and 10 deletions per sample using the X-replacement method in each of the four datasets. As the X-replacement method was employed, deletions were simulated only in the X chromosome of female samples from each respective data set. The number of simulated CNVs ranged from 72 - 144 for each dataset (see supplementary material). The simulated deletions spanned between 100bp and 6.9kbp of targeted bases, equating to genomic spans of between 471bp and 4.3Mbp.

**Table 2:** Estimated, actual and improved sensitivity for validated CNVs from Krumm et al. (2015). The predicted sensitivities reflect the actual sensitivity well for all callers exception XHMM.

| Caller | Predicted Sensitivity | Actual Sensitivity | Improved Sensitivity |
| --- | --- | --- | --- |
| ExomeDepth | 88% | 90% | 90% (0%) |
| XHMM | 82% | 48% | 76% (+28%) |
| Conifer | 57% | 40% | 48% (+8%) |
| cn.MOPS | 24% | 16% | 12% (-3%) |

### Comparison of CNV Detection Methods with Default Settings

First we used Ximmer to compare the accuracy of the four different CNV detection methods. Parameters for each tool were set to their defaults, except for cases where the tool setting was clearly misaligned to the simulated data. Specifically, the cn.MOPs minimum CNV width was lowered to 1, and XHMM mean number of targets were lowered to 3 to better match the generally smaller size of deletions included in the simulation.

In the Nimblegen data set, we observe that there were significant differences between the performance of the different CNV callers (Figure 3). ExomeDepth achieved substantially better absolute sensitivity than any other tool, finding 88% of all the simulated deletions compared to XHMM finding 57%. However, the precision of ExomeDepth was poor (54%) compared to XHMM (93%). A substantial difference in precision persisted even when ExomeDepth results were filtered to equivalent sensitivity as XHMM. Therefore in this case the optimal caller is a trade off between sensitivity and specificity. Both cn.MOPs and Conifer performed poorly in terms of sensitivity, each finding less than 30% of simulated deletions. cn.MOPs has very poor precision in this data set (0.5%), and appears to output many very high confidence calls that are ranked higher than the true positives it detects.

**Figure 3:**
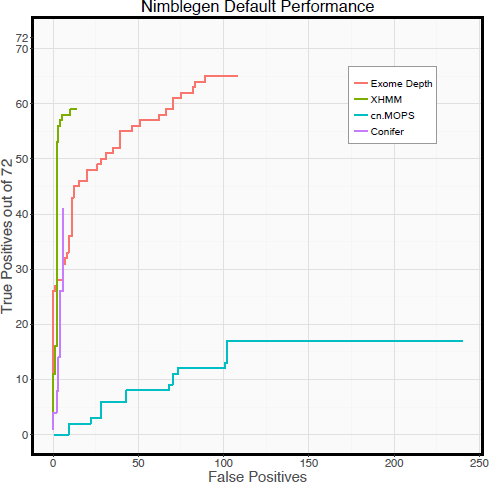
Performance of CNV callers on Nimblegen data with default settings. Performance differs greatly between callers. ExomeDepth has significantly higher sensitivity than any other caller, while Conifer and XHMM have significantly better precision.

## Comparison between Data Sets

We next compared Ximmer results using the four CNV callers with default settings on all four data sets. Our results (Figure 4) show that individual methods have marked differences in performance on different data sets. For example, all callers exhibited a low false positive rate when applied to the SureSelect data (fewer than 10 false positive calls for any caller), but showed much higher false positive rates on Nextera data (ExomeDepth and cn.MOPs both having more than 200 false positive calls). cn.MOPs performed poorly on the SureSelect, Nimblegen and Nextera data, detecting very few true and many false CNVs. However cn.MOPs is arguably the best caller for the TruSeq data, having a lower false positive rate than ExomeDepth and Conifer, but higher sensitivity than XHMM. These differences suggest that some data sets are better suited to the algorithms or default settings of particular calling methods.

**Figure 4:**
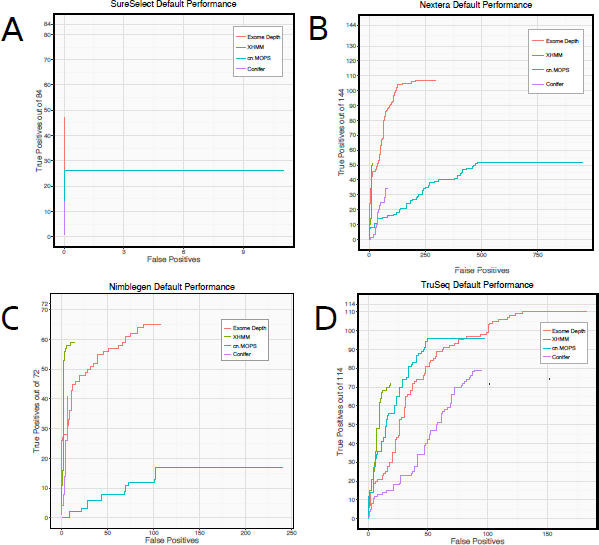
ROC-style Curve with Default Parameters - Count of true positives vs false positives as ranked results are filtered by varying quality score threshold, when the four CNV calling methods are executed on four different data sets with their default parameters. Performance is highly variable both between different methods on the same data set, and between the same method on different data sets

Despite the differences, some aspects of individual caller performance were consistent across all datasets: XHMM consistently achieved the lowest false positive rate for a given sensitivity, and conversely ExomeDepth consistently achieved higher total sensitivity than any other caller. In some respects differences between datasets are consistent between callers. With SureSelect data (30x mean coverage), no caller could achieve more than 60% sensitivity. However with TruSeq data (90x mean coverage), all callers found more than 60% of the simulated deletions, and ExomeDepth found nearly all deletions (96%).

It is likely that homogeneity of the data is an important factor in determining these characteristics: data sets having very low inter-sample variation with few significant batch effects may work well with callers that apply relatively little normalisation or are flexible in their normalisation approach.

Overall our results suggest that each data set has individual characteristics that affect the performance of each CNV caller differently. Consequently, there is no single best CNV detection tool for all data sets. Depending on the priorities of the investigation, and the particular data set in question, a different tool or combination of tools may be more appropriate. Therefore users should assess their own data and choose CNV detection methods using Ximmer.

## CNV Calling performance can be improved with parameter optimisation

We next configured Ximmer to re-analyse the Nimblegen data while varying several configurable parameters of each CNV caller. The parameters varied were chosen by reviewing the documentation and experimenting to find those having the largest direct effect on sensitivity (Table S2).

We found that adjusting two parameters (the exome-wide CNV rate to 10-4 and the normalisation factor to 0.2), increased XHMM sensitivity (Figure 4A) by 21% (67% to 88%) with an acceptable loss of precision (81% to 55%). Similarly, we evaluated alternative values for the SVD number and calling threshold for Conifer (Figure 4C), and found that, by adjusting the calling threshold parameter from 1.5 down to 1.25, sensitivity could be improved from 25% to 40% with only a small sacrifice in precision. cn.MOPs adjustments were able to improve sensitivity from 19% to 36% by adjusting the prior impact parameter from 5 to 2, and the calling threshold upwards from −0.8 to −0.4. Although we tried varying two parameters (transition probability and expected CNV length), ExomeDepth appeared to have nearly optimal parameters as its defaults for this data set.

**Figure 4:**
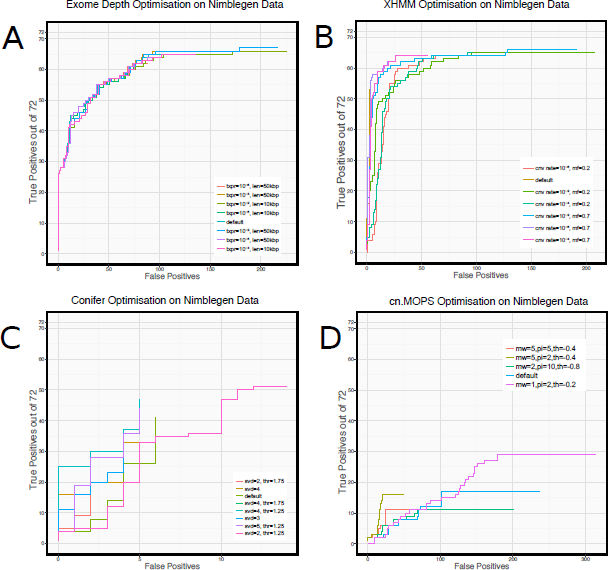
Results of adjusting CNV calling parameters on ROC-style curves. XHMM, Conifer and cn.MOPs all have configurations where sensitivity or precision can be substantially improved: reducing Conifer calling threshold to 1.25 increases sensitivity from 25% to 40%; increasing the exome-wide CNV rate to 10^−4^ and reducing the normalisation factor from 0.7 to 0.2 increases XHMM sensitivity from 67% to 88%; Reducing the cn.MOPs prior impact factor to 2 and raising the calling threshold to −0.4 allowed sensitivity to nearly double (from 24% to 36%), however these settings caused a substantial reduction in precision.

This analysis demonstrates that tuning parameter settings should be considered an important element of usage of CNV detection tools, and can lead to significantly improved accuracy. Many previous comparison studies [17, 18] have been evaluated without rigorous optimisation of parameters. Our results suggest that the discrepancies in the results from these studies may have been reduced if calling parameters were optimised.

## Optimisation of Parameters across Data Sets

We applied the optimised settings derived from simulation performance on Nimblegen data to the analysis of the other three data sets. However, we observed that these settings are not optimal for every other data set. For example, on the SureSelect and TruSeq data (Figure 5A, 5B), XHMM achieves both high sensitivity (85%) and precision (80%) with the default settings, but produces no calls at all with the optimised settings. The optimisations increase sensitivity in cn.MOPs, however, the marginal increase (69% to 77%) is much less significant than for Nimblegen data, and causes a substantially higher number of false positive calls. Similarly Conifer also shows a much smaller proportional increase in sensitivity (64% to 73%) and experiences a significant fall in precision (75% to 59%).

**Figure 5:**
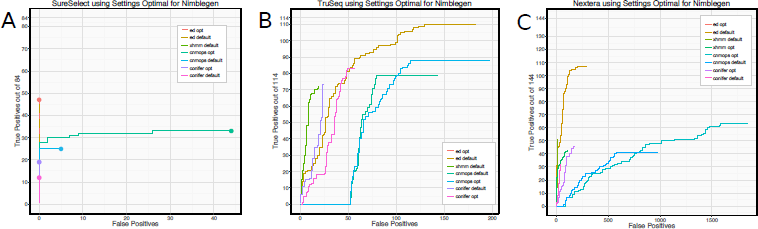
Performance of other data sets (A: SureSelect, B: TruSeq, C: Nextera) when analysed using parameters optimal for Nimblegen data (opt), compared to default settings (default). Nimblegen-optimised parameters are frequently unsuitable on other datasets. XHMM is severely compromised by the Nimblegen settings on all datasets: SureSelect and TruSeq data, produce no XHMM CNV calls, while both sensitivity and precision are poorer in Nextera data. Conifer and cn.MOPs both gain in sensitivity, but by a much smaller proportion and with a larger inflation of false positive calls than with Nimblegen data.

We conclude that optimisation needs to be performed on each data set or data type separately. Ximmer supports this process efficiently and easily through modifying simple configuration settings.

## Application of Ximmer to a set of Validated CNVs

We extracted a set of validated CNVs for the samples that were captured in the Nimblegen data set from a previous study by N. Krumm et al. (2015). After filtering to include only CNVs overlapping autosomal target regions of the exome capture, 25 validated CNVs remained. We then tested detection of these CNVs from the exome data, first using default parameters as described above for each of the four CNV callers. With the exception of XHMM, the sensitivity estimated by Ximmer using simulation approximately reflected the sensitivity observed on the validated CNVs (Table 2). In the case of XHMM we suspect that differences in the composition of the CNV sizes between the simulation and the validated CNV set may partially account for the discrepancy. Precision is harder to evaluate as predictions of CNVs in regions not in our validated set could be true deletions. However, the number of total detections varied greatly between callers (Conifer and XHMM having fewer than 12, compared to ExomeDepth and cn.MOPS having more than 200), as predicted by Ximmer.

We next applied the optimised settings identified previously through simulation to improve sensitivity for ExomeDepth, XHMM and Conifer. Due to the poor precision observed with the default settings for cn.MOPS, we chose to improve precision rather than sensitivity. By reviewing the sample counts plot, we identified that a significant fraction of the putative false positive calls were concentrated in just 3 out of 20 samples (Supplementary Figure S7). Therefore we excluded these samples from the analysis for cn.MOPS.

The results after incorporating parameter adjustments suggested by Ximmer show substantially improved performance of several methods (Figure 6). For example, sensitivity was improved in XHMM (+28%) and Conifer (+8%). Conversely, removing the three poor quality samples from cn.MOPs slightly lowered sensitivity, but removed 90% (856) of the false positive CNV calls.

**Figure 6:**
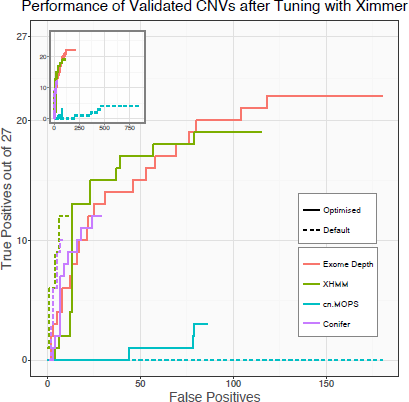
Comparison before and after applying adjusted parameters to improve performance. XHMM and Conifer sensitivity are substantially improved (+28%, +8%) with minimal loss of precision, while no improvement was possible for ExomeDepth. cn.MOPS precision was substantially improved by excluding poor quality samples identified by Ximmer’s sample counts plot

## Conclusion

While there is great utility in detecting CNVs from WES data, adoption of CNV detection methods in practice has met with significant challenges. These are primarily centred around highly unpredictable performance and lack of reproducibility between data sets. We have addressed these challenges by creating Ximmer, a tool that facilitates efficiently assessing and improving the accuracy of WES-based CNV detection methods. Our comparison of four different data sets analysed by four different CNV calling methods represents one of the most comprehensive evaluations to date. Our results show, consistent with previous studies, that there is significant variability in performance of CNV detection between tools and between data sets. We conclude that to effectively use these methods, attention must be applied to understand and optimise their behavior on each individual data set. Ximmer can be used to automate these procedures, avoiding a significant burden. In addition, we have demonstrated that Ximmer can produce valuable insights into the quality of data sets for CNV calling and the behavior of CNV detection tools. As the first tool offering combined simulation, evaluation, tuning and interpretation of results from CNV detection methods, we believe Ximmer will assist increasing practical adoption of CNV detection methods for exome data. Ximmer is open source and available at http://ximmer.org. An example Ximmer report can be viewed online at http://example.ximmer.org.

## Authors’ Contributions

SS conceived of and implemented the method, performed simulation work and drafted the manuscript. AO advised on design, implementation and interpretation of results, and edited the manuscript. SM and JE collected samples, extracted DNA, arranged sequencing and provided feedback on the manuscript. All authors approved the final manuscript.

## Additional Files

### Additional File 1

A set of figures, tables and supplementary methods supporting the results. Details of the simulation method. Description of methodology for optimising tuning parameters of data sets. **Table S1.**

Description of confidence measures used to rank CNV calls. **Table S2**. Parameters selected for optimisation of CNV detection methods. **Figures S3 – S5**. ROC-style curves for optimisation of CNV detection methods. **Figure S7**. Sample count QC plot showing frequency of CNV calls for each sample across different cn.MOPs configurations.

## Acknowledgements

We thank Harriet Dashnow for reading and providing feedback on the manuscript. We thank the Melbourne Genomics Health Alliance for access to exome sequence data from the demonstration project. This work was made possible through Victorian State Government Operational Infrastructure Support and Australian Government NHMRC IRIISS. This work was supported by the National Health and Medical Research Council, Australia (Career Development Fellowship 1051481 to AO)

Sequencing for the TruSeq data set was provided by the Center for Mendelian Genomics at the Broad Institute of MIT and Harvard and was funded by the National Human Genome Research Institute, the National Eye Institute, and the National Heart, Lung and Blood Institute grant UM1 HG008900 to Daniel MacArthur and Heidi Rehm.

